# Loss of wings induces the expression of the worker-like phenotype in queens of a ponerine ant

**DOI:** 10.1101/2021.10.16.464676

**Authors:** Benjamin Pyenson, Christopher Albin-Brooks, Corinne Burhyte, Jürgen Liebig

## Abstract

Many highly-eusocial insect species are characterized by morphological differences between females. This is especially pronounced in ants where queens usually possess a fully developed thorax with wings and are specialized for reproduction while workers have a reduced thorax without wings and show various levels of reproductive degeneration that is associated with their helper role in the colony. Despite their morphological differentiation, queens and workers still show some plasticity leading to overlapping behavioral and physiological phenotypes. We investigated the level of queen plasticity and the factor that induces a worker-like phenotype in the ant species *Harpegnathos saltator* that has limited queen-worker dimorphism and workers that can assume the reproductive role of a queen in the colony. By comparing alate and dealate young queens, so-called gynes, we found that the loss of wings initiated the expression of behavioral and physiological characteristics of ant workers. In contrast to alate gynes, dealate gynes displayed higher frequencies of worker-like behaviors. In addition, dealate gynes showed a worker-like range of reproductive states unlike alate gynes. Like workers, dealate gynes lost the chemical signaling that is characteristic of alate gynes. Since gynes can activate this worker-like phenotype after wing loss, the essential difference between the ant queens and workers in this species with limited queen-worker dimorphism is a dispersal polyphenism. If the phenotypic plasticity observed in *H. saltator* is representative of the early stages of ant eusociality, an emerging dispersal dimorphism rather than a distinct reproductive dimorphism might represent one of the first steps in ant evolution.

**Summary Statement:** Ant queens who have lost their wings express worker-like behaviors and physiology including the display of dominance behavior during hierarchy establishment, which is normally a worker-only behavior in this species.

## Introduction

Alternate behavioral and physiological phenotypes from individuals with the same genome can be a response to environmental variation. This phenotypic plasticity is an important mechanism of morphological and behavioral variation across many taxa (West-Eberhard, 1989). Alternate phenotypes, such as eyespots on wings, the presence or absence of wings, and variation in horn length, have been associated with various non-mutually exclusive mechanisms, like differences in gene expression, endocrine signaling, and appendage development (Shapiro, 1976; Wheeler and Nijhout, 1983; Crespi, 1988; Emlen, 1994; Evans and Wheeler, 1999; Abouheif and Wray, 2002). In social insects, complex endocrine and molecular mechanisms regulate the development of the alternate morphological phenotypes of queens and workers or different sizes of workers (Wheeler, 1986; Corona et al. 2016). In addition to the morphological variation, plasticity in behavioral phenotypes within one morph may lead to an interaction between morphological and behavioral variance.

Within Hymenoptera, the most distinct morphological differences between queen and workers are present in ants (Hölldolber and Wilson, 1990). While queens and workers can be distinguished by the degree to which anatomical modules are associated with each morphological phenotype (Londe et al. 2015), we can also define behavioral and physiological phenotypes as worker-like or queen-like depending on their association with the queen or with the worker morph (Smith et al. 2018). In most cases, the ant queen has a fully developed thorax with large muscles, wings to fly, and somewhat developed ovaries before dispersing from the nest (Fletcher and Blum, 1981; Vieira et al. 2011; Monnin et al. 2018; Peeters et al. 2020). After flying from the nest, the queen mates with males attracted by her sex pheromones (Ayasse et al. 2001). After finding a new habitat, she sheds her wings and lays eggs within several days (Keller and Passera, 1990). The queen cares for her first generation of brood, and, in some species, also forages to provision them with food (Cassill, 2002; Peeters and Ito, 2001). Eventually, the adult worker offspring perform most of the non-reproductive labor of an established colony, such as brood care, foraging, and defense, with thoracic muscles that favor ground-based locomotion (Hölldobler and Lumsden, 1980; Traniello, 1989; Keller et al. 2014; Walsh et al. 2018; Peeters et al. 2020). This clear-cut difference is nevertheless not universal.

In many natural contexts, the behavior and physiology of social insects does not strongly associate with the specific queen or worker morphology (Sumner et al. 2018). In ant species without queens, all tasks are divided among wingless workers (Peeters and Ito, 2001; Monnin and Peeters, 2008). In these and other species, reproductive workers signal their fecundity in similar ways to queens (Bourke, 1988; Peeters and Liebig, 2009). When workers reproduce, they may show dominance interactions (Fletcher and Ross, 1985; Oliveira and Hölldobler, 1990; Ito and Higashi, 1991; Ito, 1993; Monnin and Peeters, 1999) which are also expressed by queens in some social systems (Oliveira and Hölldobler, 1991; Medeiros et al. 1992; Kolmer and Heinze, 2000; Yamauchi et al. 2007). Furthermore, queens may be involved in nest construction (Peeters and Andersen, 1989; Murakami, 2020), and defense (Lachaud and Fresneau, 1985; Jerome et al. 1997) which are housekeeping tasks typically performed by workers. Because ant queens and workers show context-dependent degrees of association with specific behavioral and physiological phenotypes, how is the expression of these worker-like instead of queen-like phenotypes regulated in a morphological queen or worker?

Given that reproductive queens may show reduced behavioral and physiological plasticity, we wanted to investigate this question in gynes, which are ant queens who have not yet dispersed from the natal nest and mated. Queens are the only female ants capable of aerial dispersal (Peeters and Ito, 2001). Therefore, the expression of queen-like phenotypes associated with dispersal are impossible to induce in workers. After dispersal, the queen’s behavior changes as she and her new colony develop, making it challenging to assess the phenotype of a foundress independent from her colony’s size (Augustin et al. 2018). Gynes, however, are known to show worker-like behaviors like foraging (Plateaux, 1978; Della Lucia et al. 1993; Brown, 1999; Hora et al. 2005; Vieira et al. 2011; Silva Araujo et al. 2016), care for the brood (Fresneau and Dupuy, 1988; Bourke, 1991; Ito et al. 1996; Rüppel et al. 2002; Vieira et al. 2011), and nest defense (Forder and Marsh, 1989). Since artificial dealation of gynes, the experimental removal of their wings, in some species has activated the foundress behaviors of oviposition, nursing, and defense, dealation may also initiate other behaviors more typically associated with workers (Jemielity et al. 2006; Nehring et al. 2012).

With gynes of the ponerine ant species *Harpegnathos saltator*, we can test whether the attachment of wings mediates a worker-like phenotype and if it interacts with the morphological phenotype of queens. Workers and queens are morphologically distinct (Peeters et al. 2000), with gynes showing putative sex pheromones on their cuticles that disappear in foundresses and are absent in workers (Liebig et al. 2000). After the death of a foundress, workers perform dueling, dominance biting, and policing behaviors after which some workers become the new reproductive individuals, while others occupy non-reproductive roles in the colony (Peeters and Hölldobler, 1995; Sasaki et al. 2016). A dominance bite involves one biting ant grasping a recipient ant underneath in its mandible and jerking quickly towards itself, while at least two antennal boxing ants approaching and avoiding one another is a duel. Antennal boxing by one ant towards another that is unreciprocated is not a duel. This reproductive hierarchy among workers suggests that a high variance among reproductive phenotypes is a property of workers in this species (Liebig et al. 1998). In this study, we compared the expression of worker-like behavior and physiology of *H. saltator* gynes from up to three groups: alate gynes; gynes who have shed their wings; and gynes whose wings were artificially removed.

We predicted that the gynes lacking wings should duel and initiate dominance biting at higher frequencies than alates if they express a worker-like phenotype. Because gynes eclose as alates and shed their wings thereafter, we compared the worker-like behaviors of similarly-aged alate and dealate gynes. To control for the varying latency of a gyne to shed her wings (Table 1), we artificially removed the wings from gynes and compared the frequency of their worker-like behavior to alate gynes.

**Table 1.**
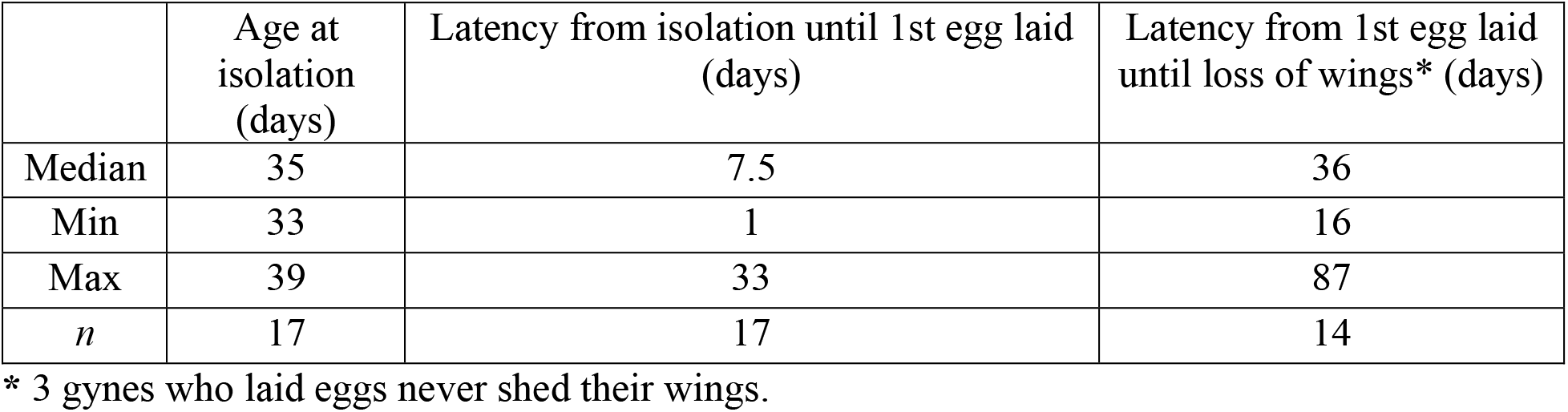
Characteristics of alate gynes in a foundress context.

|  | Age at isolation (days) | Latency from isolation until 1st egg laid (days) | Latency from 1st egg laid until loss of wings* (days) |
| --- | --- | --- | --- |
| Median | 35 | 7.5 | 36 |
| Min | 33 | 1 | 16 |
| Max | 39 | 33 | 87 |
| *n* | 17 | 17 | 14 |

\* 3 gynes who laid eggs never shed their wings.

In the presence of established reproductive individuals, we also predicted that dealate gynes, like workers, show little ovarian activity and pheromone production compared to alate gynes. We compared the reproductive physiology and pheromone production of alate and dealate gynes in the presence of an established reproductive because alate gynes may lose their wings after becoming reproductive (Table 1). Dealate gynes who shed their wings at various ages were compared to younger alate gynes. Artificially-dealate gynes were also compared to gynes of similar age who shed their wings as well as alates. Since reproductive status can require several months to manifest physiological differences in *H. saltator* workers (Ghaninia et al. 2017), we assessed these parameters two months after wing loss.

## Materials and methods

### Source Colonies

*Harpegnathos saltator* source colonies were collected from southern India from 1992 through 1999 (Peeters et al. 2000) and have been reared in the laboratory continuously since then. Individuals and brood from multiple source colonies have been bred in new combinations in the laboratory, and notes document the combination of colony backgrounds comprising a new laboratory source colony. Source colonies are distinct from one another if the combination of their originating colonies sampled from the field are different.

Colonies were maintained in a laboratory setting with a USDA APHIS permit (Number: P526P-20-01935) at Arizona State University at 25 degrees Celsius on a 12-hour photoperiod. They were housed in plastic containers (Model 79C, Pioneer Plastics, Inc. Dixon, KY, USA) lined with Fluon (Fluoropolymer resin, Dupont, Torrance, CA, USA) with a plaster base moistened with deionized water (No. 50046289, Labstone Blue, Modern Materials, Rochester, IN, USA; or No. 985-1692, Dentalstone-Buff, Darby Group Companies, Jericho, NY, USA) with an area of dugout plaster beneath a piece of glass. Two to three times a week, colonies were provided live crickets and sawdust for larvae to pupate, when frass and dead individuals were removed.

Established gamergates (mated, reproductive workers) or reproductive queens were identified in these source colonies from their reproductive behaviors and marked using Testor’s^®^ paint (Rockford, IL, USA) and/or 34-gauge artistic wire marks according to the method of Haight (2012). After laying an egg, a reproductive individual holds the egg in her mandibles and can be found near the colony’s brood, where she moves slowly compared to non-reproductive individuals. After antennal contact with a reproductive, a non-reproductive individual sometimes recoils and assume a lower submissive posture. (Liebig, 1998).

### Identification of Gynes

Pupae from source colonies were checked two to three times per week for eclosion of new gynes. Upon eclosion, new gynes were wire-marked according to the method of Haight (2012) just anterior or posterior of the petiolar node and returned to their source colony to remain non-reproductive in the presence of established reproductive individuals. Attachment of wings and mortality of these gynes was recorded when colonies were fed. In rare cases, *H. saltator* gynes show uninflated and shriveled wings after eclosion. For consistency, all gynes used in this study showed four inflated wings after eclosion. The loss of any number of wings was defined as the beginning of a dealate status. The cuticular hydrocarbons and number of yolky oocytes of a gyne were measured using procedures described below.

### Behavioral Experiment #1

To compare the worker-like dueling behavior of gynes of different alate status in the absence of established reproductive individuals, groups of 10 wire-marked gynes (age 7-40 days old, median=23 days old, *n*=20) comprised of alate gynes and dealate gynes who had already shed their wings, two or three callow workers, three worker pupae, and six male pupae were combined in a new experimental nest. One nest was created from gynes, workers, and pupae from the source colony F100, whereas another nest was created from the gynes, workers, pupae, and two males from the source colony SAFC15. Both F100 and SAFC15 showed established reproductive individuals at the time that individuals and brood were removed. Both nests were watered and fed two to three times per week and provided live crickets. Because dueling may not begin for groups of 15 individuals in a new nest until 9 days after group formation (Fig. S1), both experimental nests were not first observed until after 10 (F100) or 11 days (SAFC15). Those who had lost any of their 4 wings by the start of the analysis period were classified as dealate (age at dealation: 4-33 days, median: 12 days, *n*=7 gynes).

Gynes who participated in at least one duel with either another gyne or a worker were noted during scan observation periods of 10 minutes at least one day after feeding. Mortality and the alate status of gynes was also recorded during these periods. Alate gynes were observed 1-16 times (median: 10 observations, *n*=10 gynes) before they died or shed their wings, while dealate gynes were typically observed for more periods. Therefore, the number of periods where dealate gynes showed dueling was analyzed over only the first 10 observations to conservatively compare to the number of variable observations of alate gynes in the same nests. The gynes compared from both groups were of similar age (median= 23 days old, *n*=10 alate gynes; median=24 days old, *n*=7 dealate gynes).

### Behavioral Experiment #2

To control for the variance in the latency for gynes to shed their wings while determining whether alate status influences the expression of worker-like behavior, five randomly-chosen wire-marked gynes, one from each of five age-sorted pairs (0-26 days since eclosion, median: 10 days old, n= 110 gynes) from the same source colony, were placed into to a new plaster nest for one of two treatments: i. artificial dealation: the day of group formation, the wings of gynes were removed with Vannas Spring Scissors (3mm cutting edge, No. 15000-00, Fine Science Tools, Foster City, CA, USA) while they were held with forceps without anesthesia; ii. maintenance of alation: alate gynes were held similarly with forceps without anesthesia, but wings were not removed. 10 foraging workers from the same source colony were added to each nest.

A pair of experimental nests, one from each treatment, were filmed at 30 frames per second at 1920 × 1080 resolution with a video camera (Panasonic, model HC-VX981 or HC-V520, Newark, NJ, USA) under constant lighting but with minimal glare for 14 days continuously. Two to three times per week, all nests were fed crickets that had been previously paralyzed by workers and the nest’s plaster nest was saturated with deionized water as needed. Mortality and wing loss by alate gynes was recorded every day during filming as well as two to three times per week in the weeks after filming. The first alate gyne shed her wings 12 days after filming, while the second alate gyne lost her wings after 14 days. One alate gyne died after two days of filming, another died after 9 days, whereas one artificially-dealate gyne died four days after filming. Two to four weeks after the onset of filming, two additional gynes died in each treatment. To maximize the number of gynes showing behavior, the behavior from nests was compared within two weeks from the start of filming.

The behavior of gynes in experiment nests was analyzed ten days after the start of filming because both alate and dealate gynes will have begun to express worker-like behaviors. Experimental nests were reviewed on a computer using QuickTime as fast as 30 times normal speed from the beginning of filming to detect the first worker-like behaviors involving gynes. Workers were involved in dueling and dominance in all nests (n=6), and artificially-dealate gynes initiated dominance or were involved in dueling within six days since the start of filming in their nests (n=3). Alate gynes in two of the three nests began dueling and/or biting behavior within 9 days since the start of filming and continued to show these behaviors after 10 days (Fig. S1).

Ten days after the start of filming, all groups were reviewed as fast as 30 times normal speed for the first detection of a duel or dominance bite involving a gyne. After detecting the first duel involving a gyne, regardless of whether the partner was another gyne or a worker, the number of gyne-involved duels in the nest was counted for a subsequent two hours. The results are presented for the count of gyne-involved duels over 30-minutes because non-parametric comparisons showed similar effects of the counts of duels over 30-minutes to those over two hours (Wilcoxon tests: 30-minutes: W = 37.5, p-value = 0.1272; 2-hour: W = 36.5, p-value = 0.1116) while a parametric model comparing counts over 2-hours was overfit. The number of dominance bites initiated by a gyne in the same nest was counted for a subsequent six hours after a gyne was observed initiating the first dominance bite after 10 days of filming. In the absence of detecting at least one of these behaviors involving gynes, the interactions in a group were observed for the entirety of a 10 to14 day period.

Two weeks after filming, nests were provided live crickets two to three times per week, at which times mortality and dealation was noted. The first offspring were male, suggesting that the gynes in these nests were likely unmated. After several weeks, some of the gynes became the established reproductive individuals in these nests. The ovary development (*n*=6) of these reproductive gynes was assessed using methods described below.

### Cuticular hydrocarbon abundance and ovarian activity of non-reproductive gynes

To determine how the presence of wings affects a gyne’s reproductive ontogeny as a foundress, we isolated gynes in a small plastic container (Model 29C, Pioneer Plastics, Inc. Dixon, KY, USA) with a blue plaster base lined with Fluon as above. The gynes were checked for the presence of brood, wing loss, and mortality during observations one to two times per week before providing them pre-stung crickets and saturating their plaster with deionized water. Those gynes that shed their wings did so after laying eggs (Table 1). Because gynes become dealate after becoming reproductive, it is difficult to interpret whether the phenotype of these foundresses represents their alate status or their reproduction. To avoid this complication, we compared the ovary development and abundance of cuticular hydrocarbons (CHC) from alate gynes and dealate gynes in colonies with established reproductive individuals.

To control for the age at which a gyne loses her wings, we also non-destructively sampled CHC abundance from gynes who remained in the nests of source colonies with established reproductives at one, two, and three months of age before culling them, dissecting their ovaries, and counting their yolky oocytes. Since callow gynes show a lower abundance of large alkadienes compared to sclerotized alate gynes (Liebig et al. 2000), we sampled gynes at one month to provide sufficient time for sclerotization and the associated changes in CHCs.

After non-destructive sampling of CHC abundance, one-month old gynes were randomly assigned to one of two treatments: i. artificial-dealation: gynes were dealated as above and returned to the source colony; ii. maintenance of alation: gynes were treated as above without wing-removal and returned to the source colony. In this way, all gynes in the artificial-dealation group lost wings at 1 month, while those in the control group remained alate. One-month old gynes designated for alate or artificially-dealate treatments did not differ in the abundance of pentatriacontadiene (C35:2) and heptatriacontadiene (C37:2) (Fig. S3). Gynes from these two treatments were sampled again for CHC abundance one and two months later (i.e. at two- and three-months old, respectively). After sampling the gynes for the third time, the ovaries of gynes were dissected and the number of yolky oocytes counted.

In some cases (n=3), control alate gynes shed their wings after the initial sampling of CHC abundance at one-month old. In other cases (n=3), control alate gynes shed their wings after the second sampling of CHC abundance at two-months old. Two months after the loss of their wings, gynes from these two groups, described as naturally-dealating gynes (NATLDEALATE), were pooled for analysis of CHC abundance as well as for the number of yolky oocytes.

In rare cases (n=2), the fibers used to sample CHC abundance were stored in a refrigerator in a plastic box after sampling. Later, the fibers were allowed to warm to room temperature over a few minutes and alkadiene abundance was analyzed in the GCMS. Specifically, the CHC abundance was sampled two months after dealation and analyzed in the GC-MS 14 days later. Visually comparing these chromatograms to others in the same treatment revealed no conspicuous difference due to storage in the refrigerator.

### Sampling CHC abundance

Gynes were immobilized without anesthesia in a paper cutout such that only its gaster remained exposed on one side of the paper. To collect CHCs, a solid-phase microextraction (SPME) fiber (Supelco, 30um, PDMS, fused silica, 23-gauge, Yellow, 57289-U, Bellefonte, PA, USA) was rubbed across the gaster of the gyne’s cuticle for 500 strokes. This method has been shown to have similar effectiveness to hexane-extraction for detection of larger hydrocarbons (Moneti et al. 1997).

Except for the rare cases (n=2) where fibers were stored in a refrigerator (see above), the fiber was inserted immediately after sampling into a 280 degree Celsius inlet of a Gas-Chromatograph/Mass-Spectrometer (GC-MS), and analyzed using helium as a carrier gas, through a non-polar column (Agilent J &W, GC column, DB-1MS, size 30.0m X 250um X 0.25um nominal, Santa Clara, CA, USA), to ionization by a filament, and to the Mass-Spectrometer detector. The oven of the GCMS was programmed to begin at 60 degrees for 2 minutes, increasing by 40 degrees per minute to 200 degrees Celsius. At 200 degrees Celsius, the oven increased by only 5 degrees per minute Celsius to 320 degrees Celsius over 15 minutes. C35:2 and C37:2 eluted near the known elution times of the straight-chain alkane standards for pentatriacontane (C35) and heptatriacontane (C37) resuspended in pentane solvent that were subject to the same temperature program.

### Analyzing CHC abundance

By manually analyzing the ion fragmentation patterns of these candidate elution peaks, the alkadiene can be identified with a base ion with a molecular weight of 67 (Kroiss et al. 2011) as well as the molecular ion of 488 (C35:2) or 516 (C37:2). Once the alkadienes were identified in the chromatogram, integration of the area under all elution peaks in the chromatogram was used to calculate the peak areas of C35:2 and C37:2 as a proportion of the total peak area of all elution peaks in the chromatogram. Peak area for a given compound relative to all compounds later than the retention time of C23 straight-chain alkane in the chromatogram was used as the response variable because the total amount of hydrocarbons sampled from one gyne to the next likely varies. Nonpolar compounds that elute earlier than C23 are not present on the *H. saltator* cuticle in significant quantities (Liebig et al. 2000).

### Ovary Dissection and Yolky Oocyte Count

Gasters were separated from the rest of the body of a gyne and tergites of cuticle were separated from internal tissue in an anterior to posterior order in deionized water using forceps on a Sylgard medium (Dow, Inc., Midland, MI, USA). The ovary was then isolated from connected tissues to reveal exposed ovarioles and spermatheca. Mating status was confirmed by identifying a white opaque in the center of the spermatheca (Peeters et al. 2000). Conspicuous yellow bodies in the posterior portions of ovarioles indicated prior egg laying activity (Peeters and Hölldober, 1995). Consistent with the literature (Peeters et al. 2000), “larger yolky oocytes” was used as a proxy for ovarian activity, which could be distinguished from developing oocytes and trophocytes by their: 1) opacity indicative of the presence of yolk protein and 2) ovoid shape indicative of the chorion. Photographs of dissected ovaries and spermatheca were captured using SPOT Insight QE (Model 4.2, Diagnostic Instruments, Inc. Sterling Heights, MI, USA) on a Windows-based computer mounted on the trinocular port of a stereoscope (Leica MZ-125, Buffalo Grove, IL, USA), while the stage was illuminated by dual-fiber gooseneck (NCL 150, Volpi Mfg. Auburn, NY, USA). An observer blind to the true identities of the gynes counted the number of yolky oocytes, the presence of yellow bodies, and the mating status of the gyne from these photographs. The blind estimates of mating showed a perfect correlation with estimates freshly after dissecting, while estimates of yolky oocytes were highly correlated with those estimated by CAB (Fig. S2). All alate gynes (n=12) and most dealate gynes (n=12 of 13 artificially-dealate; n=4 of 5 naturally-dealate) were unmated based on the absence of a filled spermatheca.

### Statistical Analyses

To compare alate gynes and dealate gynes who shed their wings in source colonies, Wilcoxon’s non-parametric test was used in the analysis of the number of yolky oocytes, the abundance of cuticular hydrocarbons and age. Since gynes are produced from a limited number of source colonies, however, the response variables for experimental data were analyzed in a linear mixed-model (LMM). In these cases, treatment was the fixed effect, while the source colony identity was the random-intercept effect. To determine whether a mixed model will be a better fit than a random-effects model, Akaike’s Information Criterion was calculated and the model structure with the lowest value was analyzed. An F-test with Kenward-Roger approximation for smaller sample sizes (Kenward and Roger, 1997) was used to compare the full model to a random-effects model to indicate the significance of the fixed effect. Significance was assessed at an alpha of 0.05. All mixed models were fitted with the function *lmer* (Bates et al. 2015), and all statistics were conducted in R*(v.3.2.3).

To check the assumptions of the mixed models, Pearson’s residuals were checked for normality with a Shapiro-Wilk test at an alpha of 0.01. For those with p-value between 0.05 and 0.01, linearity of residuals was visually evaluated with quantile-quantile plots. The homogeneity of variance of Pearson’s residuals were checked visually by plotting the residuals against fitted values (Zuur et al 2009). In cases where models did not satisfy assumptions of normality, count data was ln (Y+1) transformed, while proportion data was arc-sine-square root transformed. For posthoc comparison of a fixed effect with more than two levels, estimated marginal means (package: emmeans) evaluated which groups were significantly different using Tukey adjustment of p-values for multiple comparisons.

## Results

### Behavior

If dealation activates the worker-like phenotype, dealate gynes should be involved in worker-like behaviors more frequently than alate gynes of similar age. From observations of experimental nests with dealate gynes who shed their wings at various ages (range: 4-33 days; median: 12 days; n=7) as well as similarly-aged alate gynes from the same source colony, the dealate gynes were found to be dueling more often (**Fig. 1**). To control for the latency until gynes become dealate, we removed the wings from groups of five gynes on the same day and combined them with non-reproductive workers from the same source colony in a new nest in a second experiment. These artificially-dealate gynes engaged in duels and initiated dominance bites at a much higher level than alate gynes in similar groups (**Fig. 2**).

**Fig. 1.**
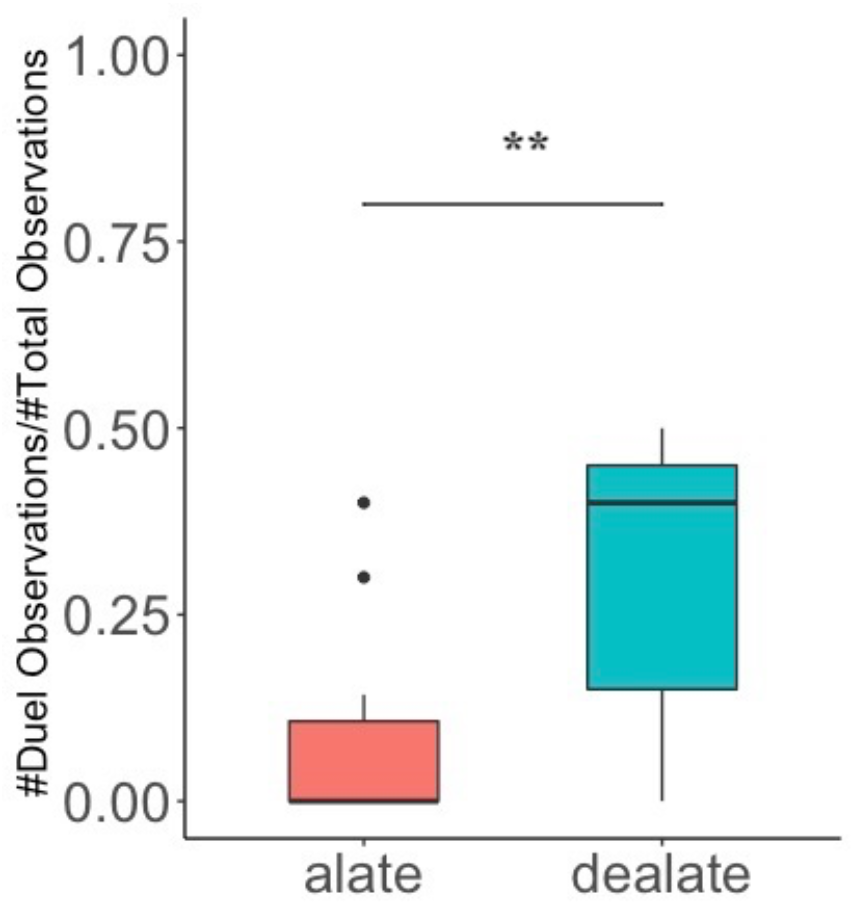
Dueling by alate and dealate gynes. Dealate gynes were observed to be dueling over a larger proportion of 10-minute scan observations than alate gynes of similar age. Box shows median and interquartile range, while whiskers extend to 95% of data, and points are outliers. **: p-value< 0.01.LMM of arc-sine-square-root-transformed data with Kenward-Roger approximation for small samples (alate, *n=*10; dealate, *n=* 7; F_1, 14.03_= 9.36, p=0.008).

**Fig. 2.**
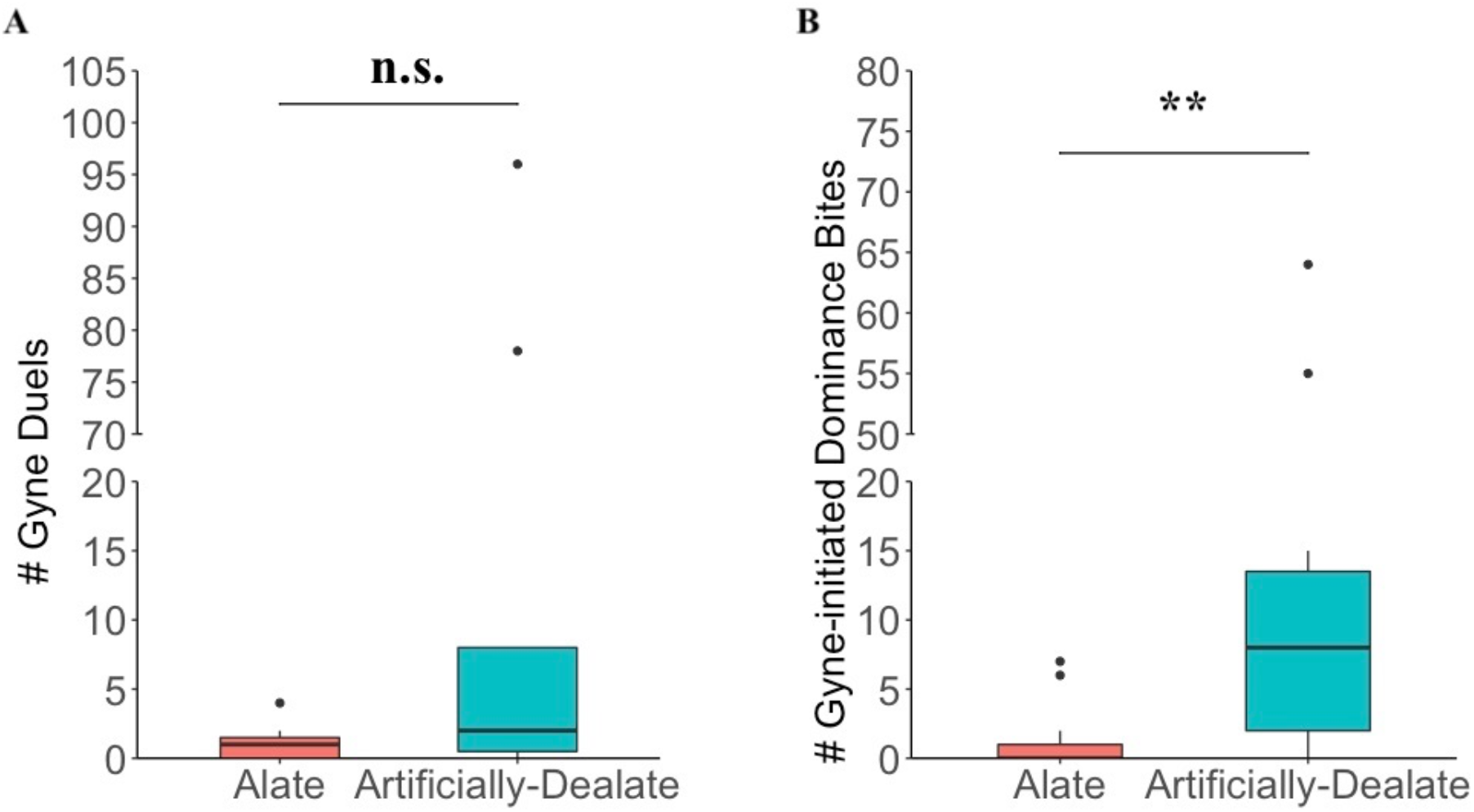
Frequency of duels and dominance bites involving gynes in the alate and artificially-dealate experimental nests. Gynes in the artificially-dealate groups are involved in more duels than the alate gynes in the alate groups. **B)** Gynes in the artificially-dealate groups initiate significantly more dominance bites. Box shows median and interquartile range, while whiskers extend to 95% of data, and points are outliers. **: p-value< 0.01. LMMs on ln (Y+1)-transformed data using F-test with Kenward-Roger approximation. (Alate groups, *n=*11; Artificially-Dealate groups, *n=* 11**; A)** F_1, 16.05_=4.05, p= 0.061; B) F_1, 16.04_=10.66, p= 0.005.)

### Ovarian Activity

Alate queens showed yolky oocytes in the presence of established reproductives. If this is associated with dispersal and dealation produces a worker-like reproductive state, then dealate gynes in the presence of established reproductive individuals should show fewer yolky oocytes than alate gynes. In source colonies with established reproductive individuals, gynes who dealated at various ages (range: 53-165 days; median: 66 days, n=7) showed fewer yolky oocytes (Alate, median: 5 yolky oocytes, *n*=8, Dealate, median: 0 yolky oocytes, *n*= 7) and tended to be older than alate gynes (**Fig. 3**). To control for the variance in age and latency of a gyne to lose her wings present in the previous comparison, we removed the wings from one-month old gynes (Artificially-Dealate) and compared their subsequent ovary development to that of alate gynes of similar age in source colonies with established reproductive individuals. To account for the effect of removing their wings, we also compared the ovaries of Artificially-Dealate gynes to those of gynes who shed their wings at various ages (Naturally-Dealate). Two months after wing loss, both Artificially-Dealate and Naturally-Dealate gynes showed fewer yolky oocytes in their ovaries than three-month old alate gynes (**Fig. 4**). While the ovaries of these alate gynes were active, they were still not as developed as those of older reproductive dealate gynes used in the second behavioral experiment (n=6, age: 377-524 days, median: 497.5 days, 7-17 yolky oocytes, median: 11.5 yolky oocytes).

**Fig. 3.**
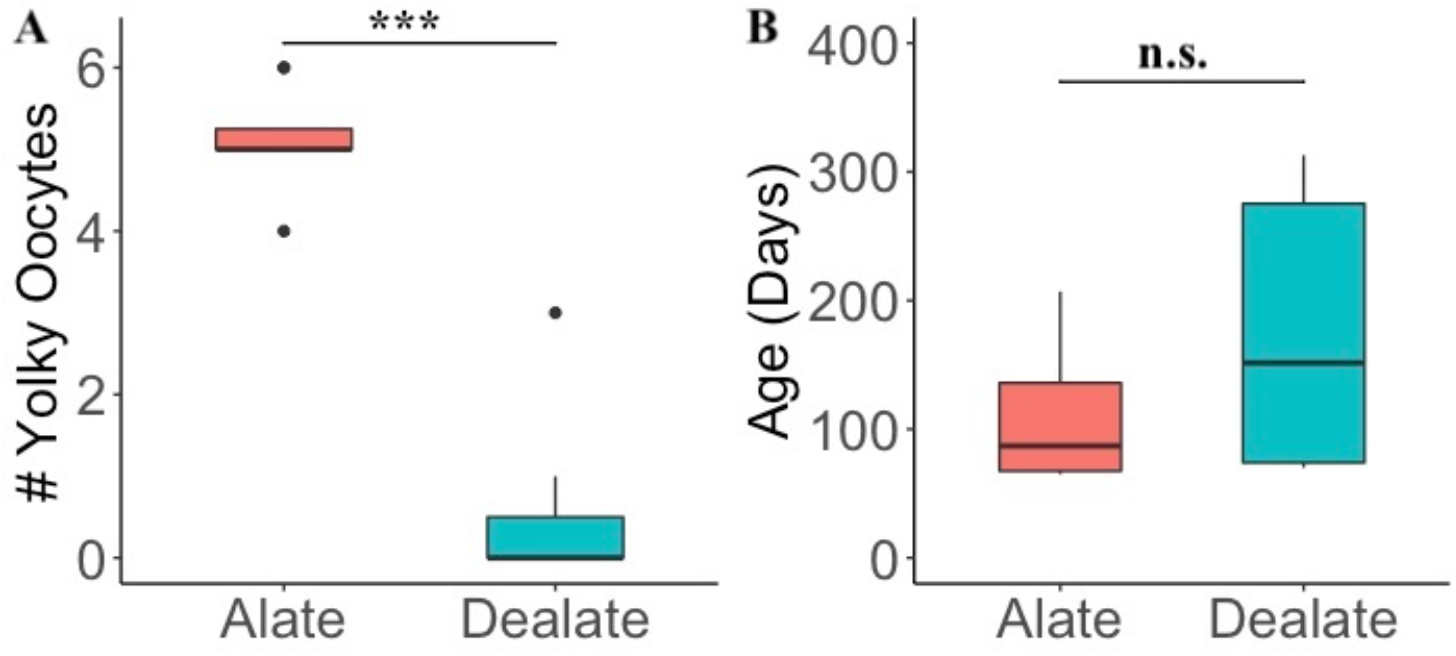
Ovarian activity of gynes of different alate status and age in colonies with established reproductive females. A) alate gynes show more yolky oocytes as counted from photographs of their dissected ovaries than dealate gynes. B) The alate gynes tend to be younger than the dealate gynes. ***: p-value< 0.001. Box shows median and interquartile range, while whiskers extend to 95% of data, and points are outliers (Alate gynes, *n=*8; Dealate gynes, *n=* 7; Wilcoxon tests: A) W = 56, p = 0.0009; B) W = 59, p = 0.076).

**Fig. 4.**
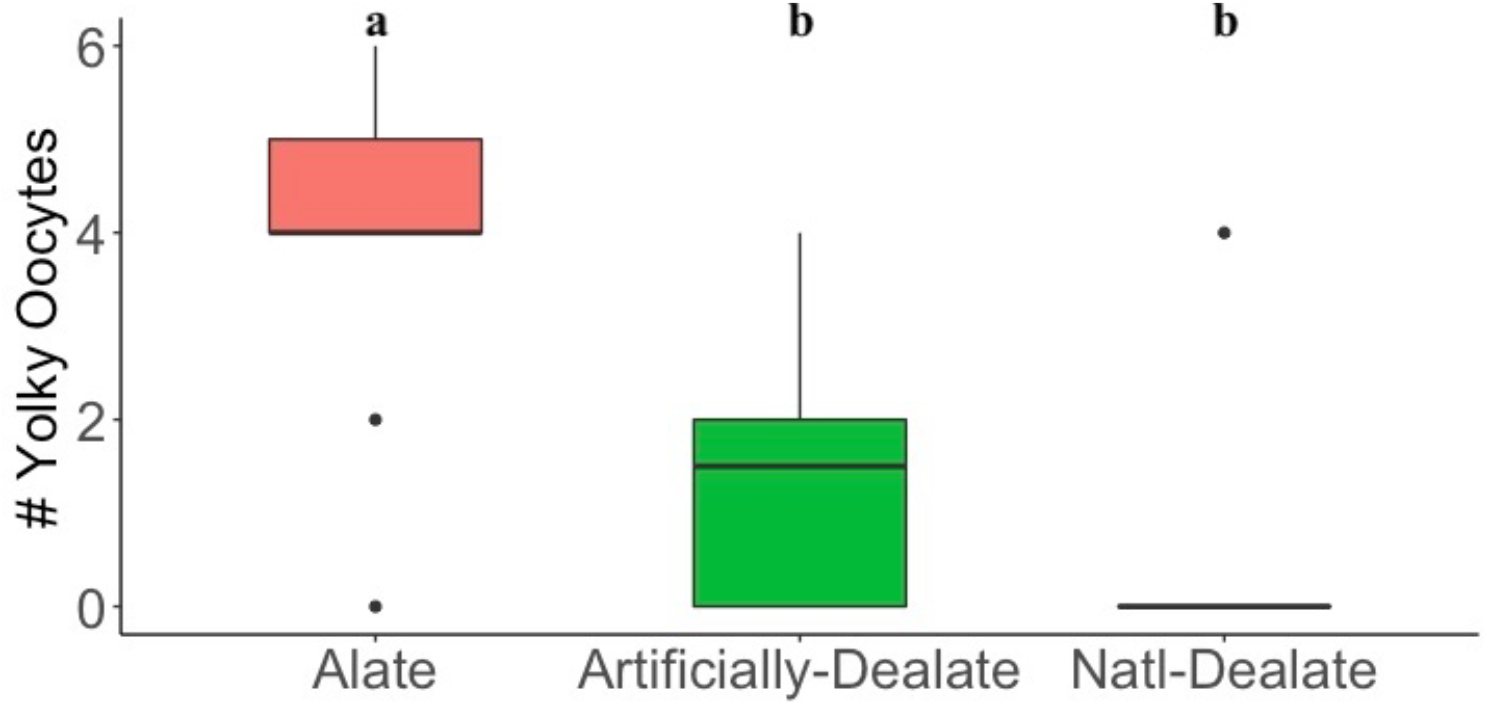
Ovary development of non-reproductive gynes depends on alate status. 2 months after wing loss, the ovaries of dealate gynes have fewer yolky oocytes than 3-month old alate gynes. Box shows median and interquartile range, while whiskers extend to 95% of data, and points are outliers (LMM, F-test with Kenward-Roger approximation, F_2, 22.88_= 11.875, p= 0.0002901. Estimated marginal mean differences with Tukey adjustment of p-values for multiple comparisons, shown with different letters above. Alate:*n* = 12, median age: 91.5 days; Artificially-Dealate: *n* = 12, median age: 90.5 days; Natl-Dealate: *n* = 5, median age: 128.5 days).

### Cuticular Hydrocarbon Abundance

If the hydrocarbon profile on a gyne’s cuticle also depends on alate status, then the gynes that have lost their wings should show a reduced abundance of the alkadienes that are characteristic of non-callow gynes (Liebig et al. 2000). In colonies with established reproductive individuals, gynes who shed their wings at various ages (range: 16-383 days; median: 94 days, n=21) showed lower abundances of the cuticular hydrocarbons (CHC) pentatriacontadiene (C35:2) and heptatriacontadiene (C37:2) than alate gynes (**Fig. 5**). Because of the large observed latency of gynes to shed their wings, we also compared the abundance of CHCs of Artificially-Dealate gynes, Naturally-Dealate gynes, and three-month old alate gynes. Two months after wing loss, both Artificially-Dealate and Naturally-Dealate gynes showed reduced abundance of C35:2 and C37:2 compared to three-month old alate gynes (**Fig. 6**).

**Fig. 5.**
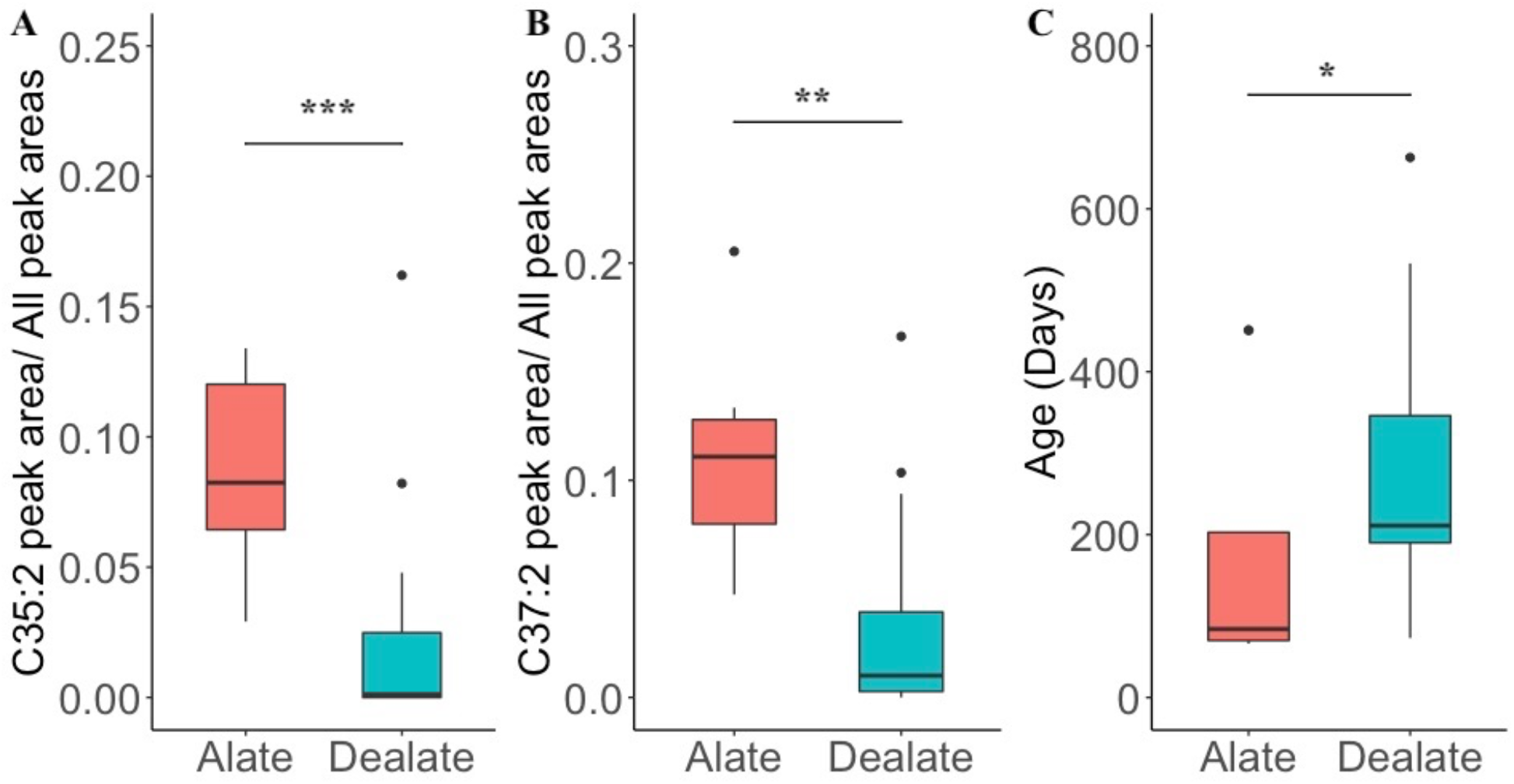
Abundance of larger alkadienes on the cuticles of gynes of different alate status and age in colonies with established reproductive individuals. The abundance of A) C35:2; B) C37:2; and C) the age of the gynes in colonies with established reproductive individuals. Box shows median and interquartile range, while whiskers extend to 95% of data, and points are outliers*:p-value<0.05; **p-value<0.01; ***: p-value< 0.001. (Alate *n*=8 ; Dealate *n*=21;Wilcoxon tests: A) W=153, ; B)W = 151, p=0.001; C)W=41.5,p=0.040).

**Fig. 6.**
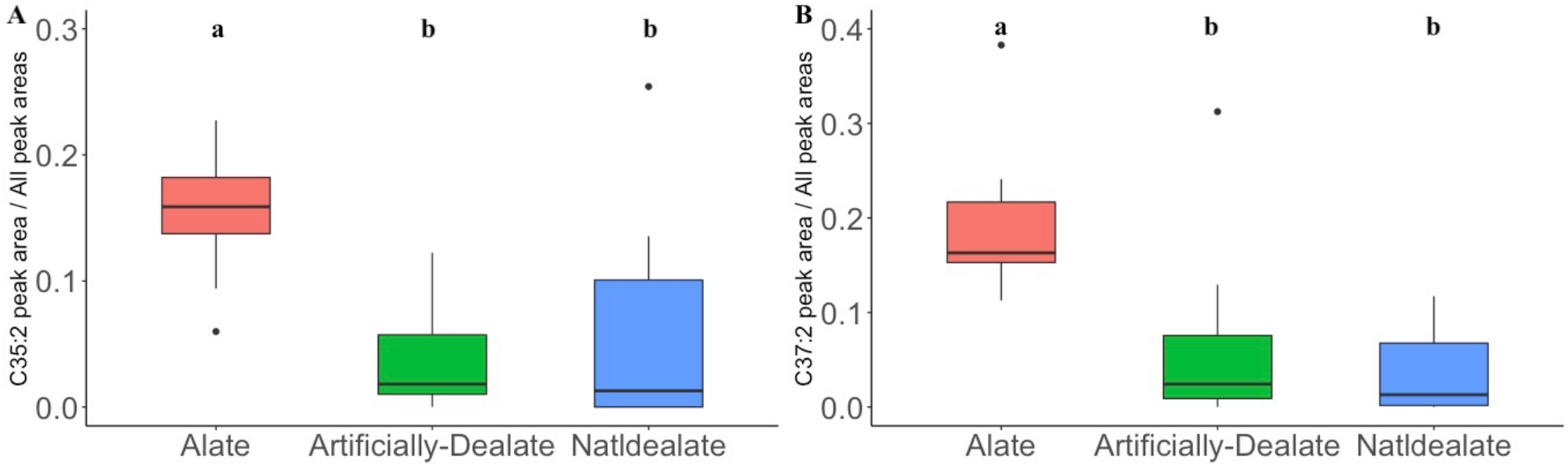
Abundance of larger alkadienes on the cuticles of gynes 2 months after wing loss or of alate gynes that are 3 months old. Alate gynes show a higher abundance of C35:2 and C37:2 than dealate gynes 2 months after wing loss. Box shows median and interquartile range, while whiskers extend to 95% of data, and points are outliers (Alate: *n* =13, median age: 91 days; Artificially-Dealate: *n* =13, median age: 90 days; NatlDealate: *n* =7, median age: 129 days). LMM of arc-sine-square root-transformed data, F-test with Kenward-Roger approximation. A) F_2, 28.59_=13.30, p=8.3e-05; B) F_2, 28.40_=14.15, p=5.5e-05. Estimated marginal mean differences with Tukey adjustment of p-values for multiple comparisons, shown with different letters above.)

## Discussion

The loss of wings of *H. saltator* gynes within an established colony induced the expression of a behavioral and physiological phenotype that is usually only shown by workers. Specifically, dealate gynes displayed behaviors that are commonly expressed by workers during the establishment of a reproductive hierarchy but not by dealate queens that follow the trajectory of founding a colony and becoming an established queen. Within a colony, alate gynes occasionally showed such behaviors as well but to a much lower extent than dealate gynes. The latter showed additional worker-like features such as low ovarian activity in the presence of reproductive individuals, but activated their ovaries in the absence of established reproductives. In contrast, alate gynes in the presence of reproductive individuals demonstrated ovarian activity associated with their preparation for a mating flight that was intermediate to non-reproductive and highly-reproductive dealate gynes. Alate gynes also expressed a higher abundance of putative sex pheromones on their cuticle that was reduced in dealate gynes as well as in workers (see also Liebig et al. 2000). These differences help identify queen-specific traits in species with little queen-worker dimorphism expected early after the transition to ant eusociality.

The presence of wings determines the expression of true worker-like or queen-like behavior in *H. saltator* gynes. In *H. saltator* and other ants, foundresses perform brood care, defense, and foraging in a laboratory setting and the field (Table 1; Peeters et al. 2000) which is also part of the task repertoire of workers (Peeters and Hölldobler, 1995; Haight, 2012). Brood care behavior can also be induced by dealation in other ants (Jemielity et al. 2006; Nehring et al. 2012). Thus, it is not clear if dealate queens express a worker-like phenotype or part of the foundress phenotype when displaying brood care behavior. In contrast, dealate gynes of *H. saltator* engaged in behaviors that are only shown by workers during the establishment of their reproductive hierarchy (Sasaki et al. 2016). Thus, the expression of these hierarchy-related worker behaviors indicate that queens switch to a true behavioral worker phenotype upon dealation within a colony. Workers on the other hand switch to a foundress phenotype that includes foraging and brood care in parallel with reproduction when isolated on their own which is a phenotype that is normally not expressed (Liebig et al. 1998). Thus, the only exclusive queen-specific phenotype is related to flying during dispersal (Hakala et al. 2019).

Wing attachment also determined the ovarian activity of a gyne. Dealate *H. saltator* gynes, like workers, showed ovarian activity that depended on social context: they were non-reproductive in the presence of reproductive inhibition; or they were highly reproductive in the absence of such inhibition (Liebig et al. 1998). However, ovaries of alate *H. saltator* gynes contained developing eggs most likely in preparation for founding a nest after the mating flight. In fact, dispersing unmated gynes of *H. saltator* have been found to have ovaries containing mature oocytes (Peeters et al. 2000). While *H. saltator* workers show different behavioral and physiological phenotypes depending on their reproductive status (Liebig et al. 1998; Penick et al. 2021), these results collectively suggest that the attachment of wings to a queen determines whether her ovarian activity oscillates as a function of social context, like workers in this species.

Unlike dealate gynes and workers, alate gynes produced a high abundance of C35:2 and C37:2 on their cuticles, which may function to attract mates (Liebig et al. 2000). Insects use sex pheromones to signal mating receptivity (Ayasse et al. 2001), but female-specific sex pheromones have only been identified from a limited number of ant species (Walter et al. 1993; Greenberg et al. 2007; Castracani et al. 2008; Greenberg et al. 2018; Iwamoto et al. 2020). While social insect queens produce various compounds that function as sex pheromones (Gary, 1962, Ayasse et al., 1999, Niehuis et al. 2013; Wen et al. 2017), sex-specific alkadienes mediate mating in some Hymenopterans (Syvertsen et al. 1995; Krokos et al. 2001) and male *H. saltator* perceive larger hydrocarbons (Ghaninia et al. 2018). Therefore, the higher abundance of alkadienes on alate *H. saltator* gynes may be a contact sex pheromone for use during the mating flight.

How might the loss of wings alter the gyne’s phenotype? While mating status and the attachment of wings influence the behavior and physiology of *Monomorium pharaonis* queens (Nagel et al. 2020), it is unlikely that mating mediates the phenotypic changes following dealation in the *H. saltator* gyne since nearly all gynes were unmated. Instead, if the wings of *H. saltator* gynes are innervated similarly to the gemmae of *Diacamma* (Gronenberg and Peeters, 1993), then these neurons may also mediate the effect of dealation on gyne behavior and physiology.

Our results indicate that queens can express the behavioral phenotype of a worker even in conditions that do normally not occur. Conversely, workers can express a queen-like phenotype as reproductive workers in a colony and in isolation similar to a founding queen (Liebig et al. 1998, Liebig et al, 2000). This means that both queen and worker morphs can express each other’s behavioral and physiological phenotype given the respective context. This also means that the main difference between queens and workers in this species is the ability of queens to disperse due to their possession of a winged thorax and flight capability. In the evolution of ants from a solitary wasp ancestor, the reproductive specialization of queens compared to workers and the loss of wings and associated change in the thoracic musculature of workers could have evolved simultaneously or one feature could have evolved first (Hanna and Abouheif, 2021). If our results apply to other species with weak queen-worker dimorphism, the difference between the queen and workers early in ant evolution might be defined by dispersal ability rather than a distinct reproductive dimorphism.

## Acknowledgments

We thank Ti Eriksson, Kevin Haight, Stephen Pratt, Jennifer Fewell, Yun Kang, and Bert Hölldobler for comments. Stephen Pratt provided storage space for videos, use of his computers for video review, and video cameras. Bert Hölldobler provided resources for crickets and the Gas Chromatograph/Mass Spectrometer.

## Supplemental Figures

**Fig. S1.**
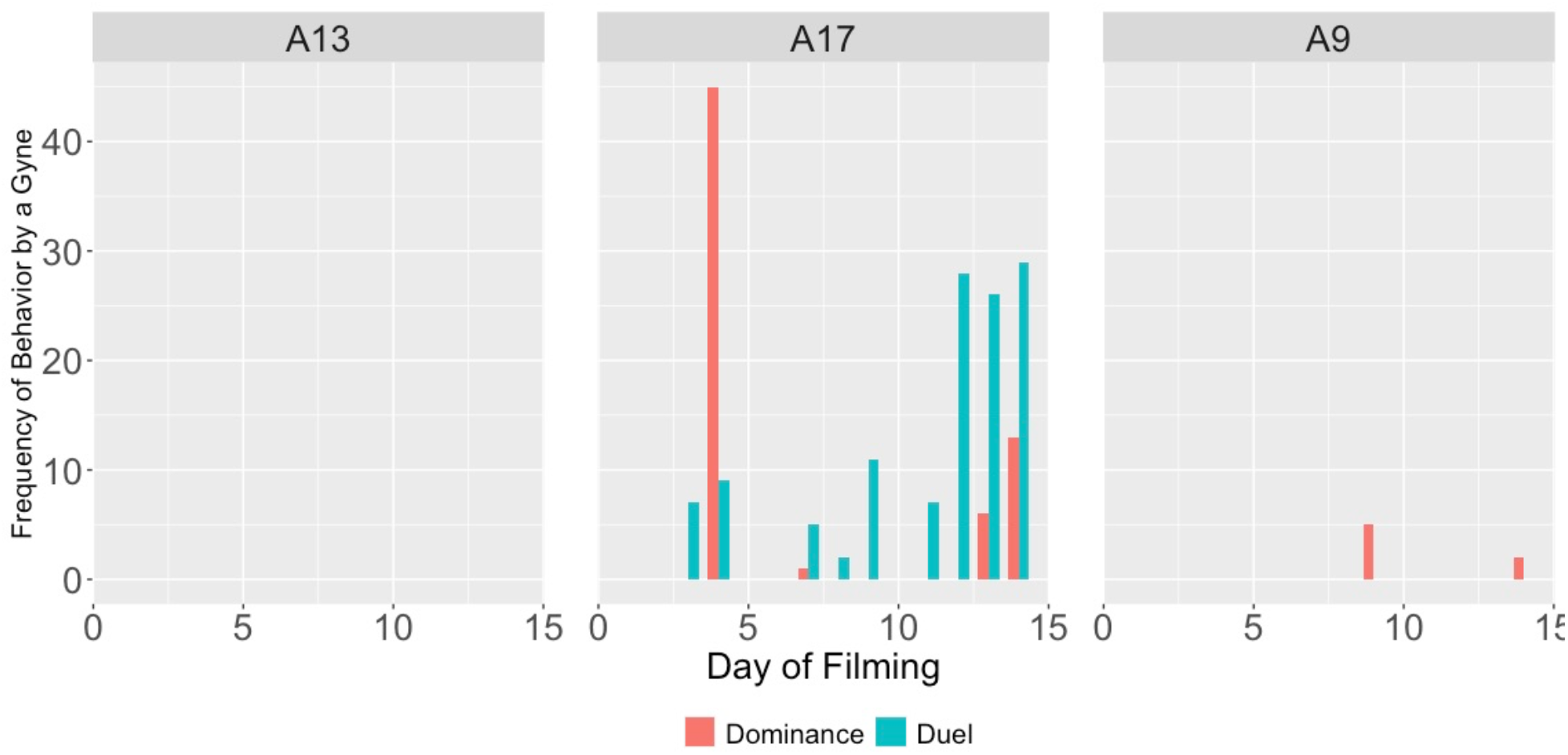
Sum of alate gyne-initiated behaviors within each day of filming over 2 weeks for 3 groups of alate gynes (A9, A13, A17) used in the second behavioral experiment.

**Fig. S2.**
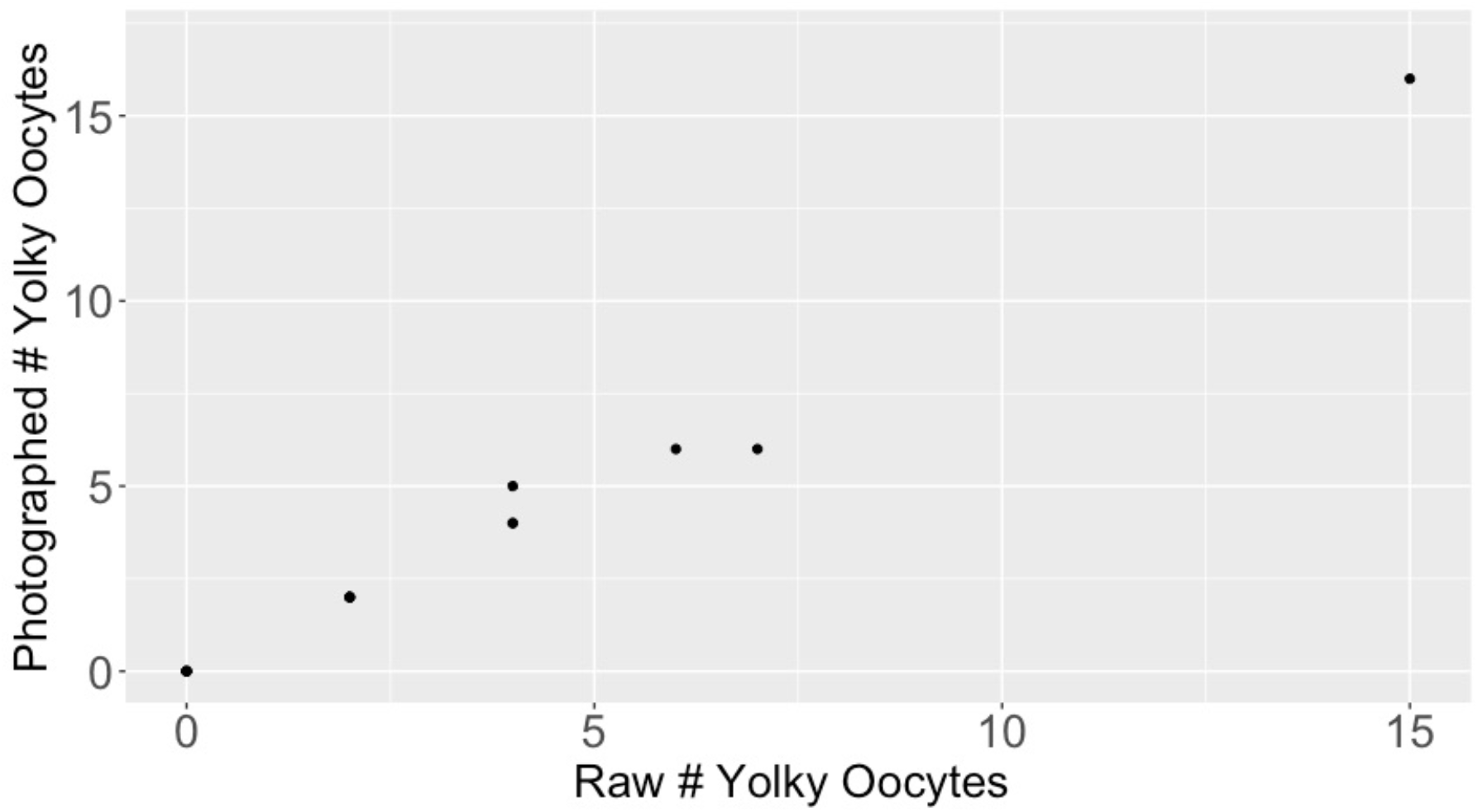
The correlation between the number of yolky oocytes of blind estimates from photographs of ovary dissections and the raw counts made by another observer of the same ovaries when they were fresh (n= 14). There is a high correlation between the number of yolky oocytes of the blind estimates from photographs of ovary dissections and the raw counts of the same ovaries when they were fresh (Pearson’s correlation: 0.99, n= 14).

**Fig. S3.**
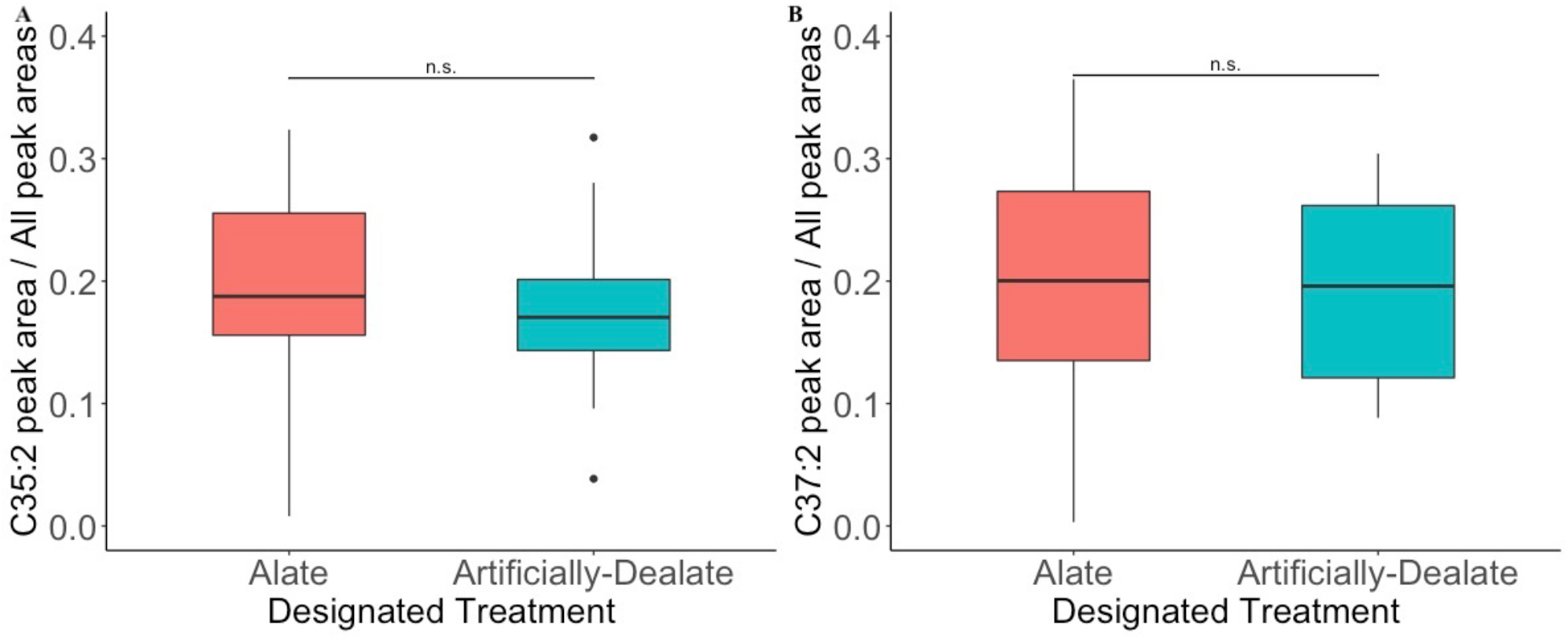
One-month old alate gynes designated for alate or artificially-dealate treatments do not differ in abundance of C35:2 or C37:2. (Alate *n=*34; Artificially-Dealate *n=* 17; Wilcoxon tests, A) C35:2, W = 306, p-value = 0.27**; B)** Wilcoxon test, C37:2, W = 264, p-value = 0.85).

